# Secreted ADAMTS-like 2 promotes myoblast differentiation by potentiating Wnt signaling

**DOI:** 10.1101/2022.12.06.519254

**Authors:** Nandaraj Taye, Mukti Singh, Clair Baldock, Dirk Hubmacher

## Abstract

The formation of multinucleated contractile myofibers from muscle stem cells during myogenesis is indispensable for skeletal muscle formation. Myogenesis is governed by myogenic regulatory transcription factors, including MYOD. However, very few MYOD-regulated effector proteins were shown to be sufficient to promote myogenesis. Here, we identified an unexpected role for the secreted matricellular protein ADAMTS-like 2 (ADAMTSL2) as a rheostat for Wnt signaling during myogenesis downstream of MYOD. ADAMTSL2 expression was induced during myoblast differentiation and ADAMTSL2 was required for myoblast differentiation. ADAMTSL2 ablation in myogenic precursor cells resulted in aberrant muscle architecture in vivo. The pro-myogenic ADAMTSL2 function was dependent on WNT ligands. Mechanistically, ADAMTSL2 potentiated WNT signaling by binding to WNT ligands and WNT receptors. Finally, we identified a WNT-binding ADAMTSL2 peptide that was sufficient to promote myogenesis. Since ADAMTSL2 was previously described as a negative regulator of TGFβ signaling in fibroblasts, ADAMTSL2 now emerges as a signaling node that could integrate and fine-tune WNT, TGFβ and potentially other signaling pathways within the dynamic microenvironment of differentiating myoblasts during skeletal muscle development and regeneration.

## Introduction

Skeletal muscles account for up to 40% of total body weight and are required for locomotion, respiration, feeding, and communication (Janssen et al. 2000). Skeletal muscles are powered by the reversible contraction of multinucleated myofibers, which are formed during embryonic and postnatal development in a process called myogenesis. During myogenesis, myogenic precursor cells differentiate into proliferating myoblasts, which exit the cell cycle and become myocytes. Myocytes then fuse with each other to form multinucleated myofibers (Petrany and Millay 2019). Myogenesis is governed by the sequential expression of the myogenic regulatory transcription factors MYF5, MYOD, MYOG, and MRF4 and supported by trophic factors secreted by muscle connective tissue cells and other muscle-resident cell types (Zammit 2017; Wosczyna et al. 2019; Theret et al. 2021). MYOD deficiency resulted in increased proliferation of myoblasts and aberrant myogenic differentiation (Guo et al. 1995; Megeney et al. 1996; Sabourin et al. 1999). Since MYOD is a transcription factor, myogenic differentiation itself is executed by MYOD-regulated effector molecules, such as secreted growth factors and their regulators or myofiber specific proteins (Bergstrom et al. 2002; Cao et al. 2010). However, very few effector proteins downstream of MYOD were shown to be sufficient to rescue lack of differentiation of MYOD-deficient myoblasts. One was MYOG, which in itself is a myogenic regulatory transcription factor (Zhang et al. 2020). In vivo, MYOD-deficiency was compensated by upregulation of MYF5 (Rudnicki et al. 1993).

Signaling pathways that regulate myogenesis include TGFβ signaling, which can inhibit myogenesis by repressing MYOD, and WNT signaling, which can promote myogenesis by inducing MYOD and other myogenic regulatory factors (Filvaroff et al. 1994; Tajbakhsh et al. 1998; Cossu and Borello 1999; Liu et al. 2001; Borello et al. 2006; Suzuki et al. 2015; Girardi and Le Grand 2018). During early development, canonical Wnt signaling mediated by WNT1, WNT3a, WNT4, and WNT6 is required for dermomyotome induction and organization as well as specification of number and type of myofibers (Linker et al. 2005; Hutcheson et al. 2009).

Non-canonical WNT signaling mediated by WNT7a (through protein kinase C, PKC) and WNT11 (through the planar cell polarity pathway) can also induce MYOD expression and regulate myocyte elongation, respectively (Brunelli et al. 2007; Gros et al. 2009). Thus, the balance and amplitude of TGFβ and canonical and non-canonical WNT signaling may ultimately influence the efficiency of myogenesis and fine-tune skeletal muscle development.

Secreted matricellular proteins are key players in the regulation and integration of signaling pathways, including WNT and TGFβ signaling, where they typically act as rheostats by providing latency to growth factors or potentiating their signaling capacity, establishing growth factor gradients, or regulating individual growth factor - receptor interactions.(Berendsen et al. 2011; Murphy-Ullrich and Sage 2014; Robertson and Rifkin 2016; Lukjanenko et al. 2019). The matricellular protein ADAMTS-like 2 (ADAMTSL2), which shares homology with ADAMTS proteases but lacks their protease domain, was identified as a negative regulator of TGFβ-beta signaling in bronchial smooth muscle cells, fibroblasts, and growth plate chondrocytes (Koo et al. 2007; Le Goff et al. 2008; Hubmacher et al. 2015; Delhon et al. 2019; Rypdal et al. 2021).

In these contexts, ADAMTSL2 repressed TGFβ signaling likely through its interaction with fibrillin microfibrils and associated proteins that regulate the bioavailability of TGFβ ligands (Le Goff et al. 2008; Hubmacher et al. 2015; Rypdal et al. 2021; Liu et al. 2022). In humans, pathogenic variants in *ADAMTSL2* cause geleophysic dysplasia (GD), which is characterized by aberrant musculoskeletal development including short stature, joint stiffness and skeletal muscle (pseudo)hypertrophy, and ∼30% juvenile lethality due to cardiac valve and bronchial and tracheal abnormalities (Le Goff et al. 2008; Marzin et al. 2020). These findings underscore an important role for ADAMTSL2 in musculoskeletal development through mechanisms that remain to be fully elucidated.

Here, we used in vitro and in vivo gain- and loss-of-function studies to interrogate the role of ADAMTSL2 in skeletal muscle development and identified an unexpected role for ADAMTSL2 as a positive regulator of myogenesis. Remarkably, knockdown of ADAMTSL2 abolished myoblast differentiation indicating an essential role in myogenesis. Indeed, ADAMTSL2 was expressed during myoblast differentiation downstream of MYOD and its overexpression promoted myoblast to myotube differentiation. Mechanistically, ADAMTSL2 potentiated WNT signaling by binding to multiple WNT ligands and the canonical WNT co-receptor LRP6, suggesting that ADAMTSL2 may increase the signaling potency of endogenous W NT ligands during muscle differentiation. Conditional deletion of *Adamtsl2* in MYF5-positive myogenic progenitor cells during development resulted in increased muscle weight, mimicking muscle manifestations in patients with GD. Collectively, these findings identify ADAMTSL2 as a rheostat for WNT signaling and a critical regulator of myogenesis downstream of MYOD.

## Results

### ADAMTSL2 is required for myoblast differentiation

We used the emergence of multinucleated myosin heavy chain (MyHC)-positive myotubes following induction of myoblast differentiation by serum-reduction as an assay to investigate the function of ADAMTSL2 during myogenesis (Fig. 1A). To validate a previous report showing induction of *Adamtsl2* expression in differentiating myoblasts, we measured the kinetics of *Adamtsl2* expression in C2C12 myoblasts up to 7 days post induction of differentiation (Koo et al. 2007). C2C12 myoblasts significantly upregulated ADAMTSL2 mRNA levels post induction of differentiation with a peak at 2 days and sustained levels until day 7 (Fig. 1B). Notably, increased ADAMTSL2 mRNA expression preceded MYH4 (MyHC) induction and myotube formation. On the protein level, ADAMTSL2 was detected in MyHC-positive myocytes and myotubes but not MyHC-negative myoblasts (Fig 1C, D). We next verified *Adamtsl2* induction in wild-type primary mouse myoblasts isolated from extensor digitorum longus (EDL), which have a differentiation kinetic similar to C2C12 myoblasts (Lagord et al. 1998). Echoing expression in C2C12 myoblasts, primary myoblasts upregulated ADAMTSL2 mRNA at 1 day and protein at 3 days post induction of differentiation, again with localization restricted to MyHC positive myocytes and myotubes (Fig. 1E-G).

**Figure 1.**
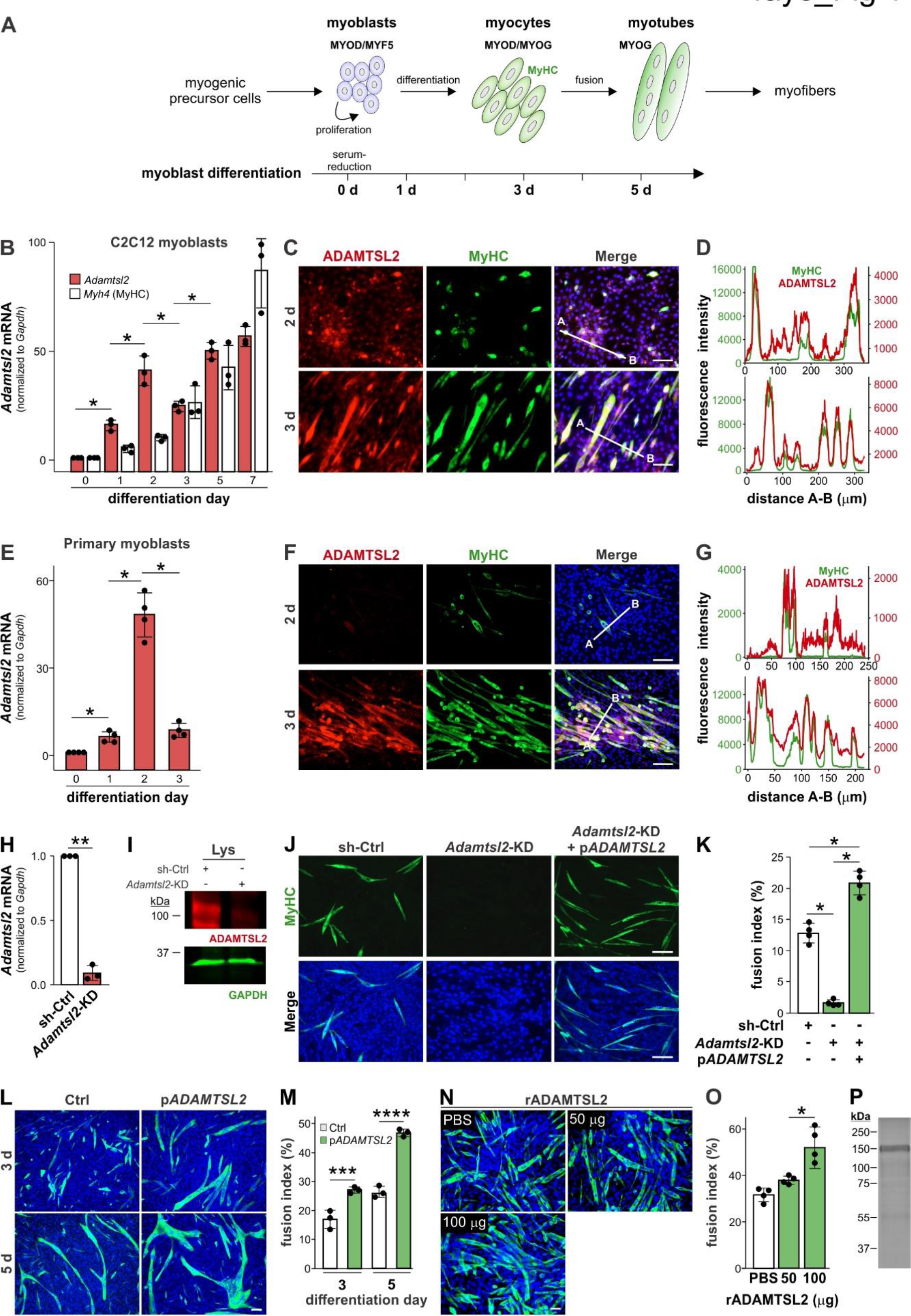
ADAMTSL2 is required for myoblast differentiation. (*A*) Diagram of myogenesis. Key stages of myoblast differentiation and associated myogenic regulatory factors are bolded. The approximate timeline of myoblast differentiation by serum withdrawal in vitro is indicated. MyHC, myosin heavy chain (MYH4). (*B*) Normalized ADAMTSL2 mRNA levels in C2C12 myoblasts undergoing differentiation by serum-reduction (n = 3). Increasing MYH4 mRNA levels indicate differentiation towards myotubes. (*C*) ADAMTSL2 and MyHC immunostaining of C2C12 myoblasts at day 2 and 3 after differentiation initiation. (*D*) Fluorescence intensity blots of green (MyHC) and red (ADAMTSL2) channels along the lines indicated in C. (*E*) Normalized ADAMTSL2 mRNA levels in primary myoblasts from wild-type EDL muscle undergoing differentiation (n = 4). (*F*) ADAMTSL2 and myosin heavy chain (MyHC) immunostaining of primary myoblasts at day 2 and 3 after differentiation initiation. (*G*) Fluorescence intensity blots of green (MyHC) and red (ADAMTSL2) channels along the lines indicated in F. (*H*) Normalized ADAMTSL2 mRNA levels in proliferating C2C12 myoblasts after stable knock-down of ADAMTSL2 with shRNA (*Adamtsl2*-KD) compared to controls (sh-Ctrl) expressing non-targeting shRNA (n = 3). (*I*) Western blot analysis of ADAMTSL2 in myoblast lysates (Lys) from *Adamtsl2*-KD and sh-Ctrl cells. (*J*) MyHC immunostaining of sh-Ctrl and *Adamtsl2*-KD myoblasts at day 3 after differentiation initiation and rescue of myotube formation by transient overexpression of ADAMTSL2 (p*ADAMTSL2*). (*K*) Quantification of fusion index from J (n = 4). (*L*) MyHC immunostaining of vector control (Ctrl) or *ADAMTSL2* overexpressing C2C12 myoblasts at day 3 and 5 after differentiation initiation. (*M*) Quantification of fusion index from L (n = 3). (*N*) MyHC staining of C2C12 myoblasts differentiated for 3 days in the presence of 0 (PBS), 50 or 100 μg recombinant ADAMTSL2 (rADAMTSL2). (*O*) Quantification of fusion index from N (n = 4). (*P*) Coomassie brilliant blue staining of 5 µg recombinant mouse ADAMTSL2 after SDS-PAGE. Scale bar: 100 µm. Bars in B, E, H, K, M, O represent mean ±SD. P-values were determined by Student’s t-test (H, M) or one-way ANOVA and posthoc Tukey test (B, E, K, O). *p<0.05, **p<0.01, ***p<0.001, ****p<0.0001.

To understand the role of ADAMTSL2 in myoblast differentiation, we depleted ADAMTSL2 mRNA in C2C12 myoblasts by stable expression of an ADAMTSL2-targeting shRNA (*Adamtsl2*-KD), which significantly reduced ADAMTSL2 mRNA and protein compared to control shRNA (sh-Ctrl) (Fig. 1H, I). Strikingly, *Adamtsl2*-KD myoblasts failed to differentiate as indicated by the absence of MyHC-positive myocytes or myotubes (Fig. 1J). As a result, the fusion index, i.e. the percentage of nuclei within multinucleated MyHC-positive myotubes, was significantly reduced (Fig. 1K). Lack of differentiation of *Adamtsl2*-KD myoblasts was fully rescued by transient overexpression of full-length ADAMTSL2 (p*ADAMTSL2*) (Fig. 1J).

Importantly, the fusion index after ADAMTSL2 overexpression was not only normalized but also increased to ∼1.6-fold compared to sh-Ctrl conditions (Fig. 1K). Differentiation of C2C12 myoblasts that constitutively overexpressed recombinant full-length ADAMTSL2 resulted in larger myotubes and the fusion index increased ∼1.8-fold (Fig. 1L, M). Finally, addition of recombinant ADAMTSL2 protein (rADAMTSL2) to C2C12 myoblasts augmented myotube formation in a dose dependent manner (Fig. 1N-P). Collectively, our data demonstrate that ADAMTSL2 is not only required for myoblast differentiation, but that its overexpression further augments myotube formation.

### ADAMTSL2 is a MYOD target gene and an effector of the MYOD-regulated myoblast differentiation program

To identify molecular pathways that could explain the lack of differentiation of *Adamtsl2-*KD myoblasts, we determined differentially expressed genes (DEGs) of *Adamtsl2-*KD compared sh-Ctrl C2C12 myoblasts at 3 days after initiation of differentiation by RNA sequencing (RNAseq). Consistent with a lack of differentiation, we observed downregulation of myogenic transcription factors (*Pax7*, *Myod1*, *Mrf4*, *Myog*), myosin heavy chain (*Myh4*) and genes required for myocyte fusion (*Mymk*, *Mymx*) (Fig. 2A). We confirmed downregulation of these genes in an independent sample set and conversely found upregulation of the same genes when ADAMTSL2 was overexpressed in C2C12 myoblasts (Fig. 2B, C). Since myoblast differentiation requires cell cycle exit, we measured cell proliferation using Ki67 immunostaining. Myoblast proliferation was significantly increased in *Adamtsl2*-KD cells and decreased in ADAMTSL2-overexpressing cells, consistent with lack or promotion of myoblast differentiation, respectively (Fig. 2D-G).

**Figure 2.**
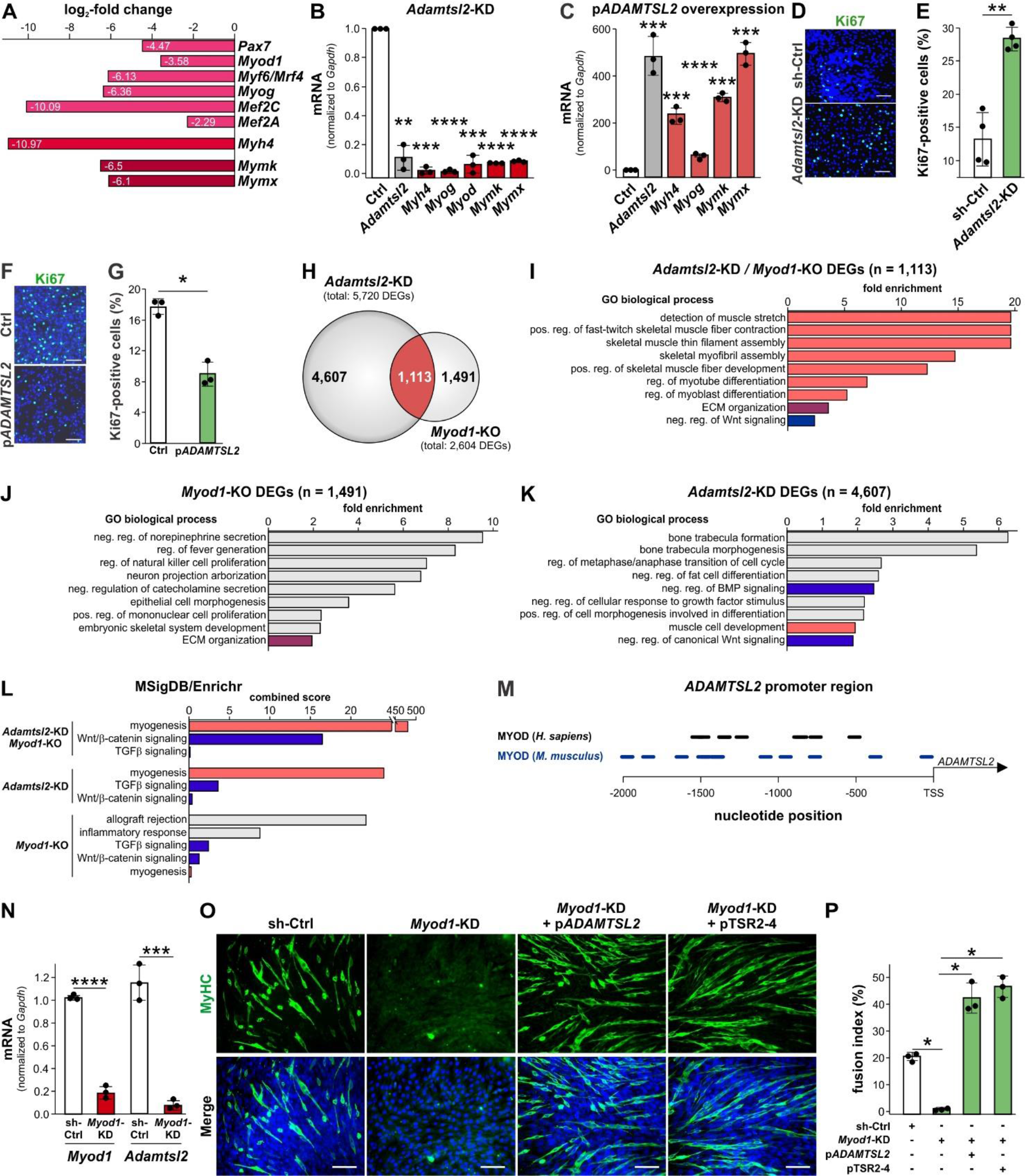
ADAMTSL2 promotes myoblast differentiation downstream of MYOD. (*A*) Log2-fold change of selected differentially expressed genes (DEGs) of *Adamtsl2*-KD compared to sh-Ctrl C2C12 myoblasts at 3 days after differentiation initiation identified by RNAseq (n=3). (*B*) Validation of DEGs from A in independent biological replicates by qRT-PCR in *Adamtsl2*-KD and sh-Ctrl C2C12 myoblasts at 3 days after differentiation initiation (n = 3). (*C*) Normalized mRNA levels in p*ADAMTSL2*-overexpressing C2C12 myoblasts at 3 days after differentiation initiation (n = 3). (*D, E*) Ki67 immunostaining for proliferating C2C12 myoblasts (D) and quantification of Ki67-positive cells (E) in sh-Ctrl and *Adamtsl2-KD* cells 3 days after differentiation initiation (n = 4). (*F, G*) Ki67 immunostaining for proliferating C2C12 myoblasts (F) and quantification of Ki67-positive cells (G) in Ctrl and p*ADAMTSL2*-overexpressing cells 3 days after differentiation initiation (n = 4). (*H*) Venn diagram of DEGs shared between *Adamtsl2*-KD and *Myod1*-KO. (*I-K*) Gene Ontology (GO) biological processes enriched in shared DEGs (I), *Myod1*-KO only DEGs (J) and *Adamtsl2*-KD only DEGs (K). (*L*) Molecular Signature Database (MSigDB) processes enriched in *Adamtsl2*-KD/*Myod1*-KO shared and distinct DEGs. (*M*) Localization of putative MYOD recognition sequences in mouse and human ADAMTSL2 promoter regions. TSS, transcriptional start site. (*N*) Normalized MYOD and ADAMTSL2 mRNA levels in *Myod1*-KD and sh-Ctrl C2C12 myoblasts (n=3). (*O*) MyHC-positive myotubes of sh-Ctrl, *Myod1*-KD, *Myod1*-KD plus overexpression of p*ADAMTSL2* or its TSR2-4 (pTSR2-4) domains in C2C12 myoblasts 3 days after initiation of differentiation. (*P*) Quantification of fusion index from O (n = 3). Scale bar in D, F, O: 100 µm. Bars in B, C, E, G, N, P represent mean values ±SD. P-values were determined by Student’s t-test (B, C, E, G, N) or one-way ANOVA and posthoc Tukey test (P). *p<0.05, **p<0.01, ***p<0.001, ****p<0.0001.

Since the *Adamtsl2*-KD phenotype is reminiscent of increased proliferation and lack of differentiation observed in MYOD-deficient myoblasts, we cross-referenced DEGs from *Adamtsl2-*KD C2C12 myoblasts with published DEGs from *Myod1* knockout (*Myod1*-KO) C2C12 myoblasts, both under differentiation conditions (Zhao et al. 2019). The Venn diagram shows that 19.4% (n = 1,113 of 5,720) of DEGs from *Adamtsl2*-KD were shared with *Myod1*- KO (Fig. 2H). Not surprisingly, DEGs that are shared between *Adamtsl2-*KD and *Myod1*-KO myoblasts were almost exclusively enriched in gene sets related to skeletal muscle, myotube formation or myoblast differentiation (Fig. 2I). Interestingly, *Myod1*-KO specific DEGs (n = 1,419 of 2,602) were enriched in neurotransmission, neuron projection arborization, immune interactions, and ECM organization (Fig. 2J). In contrast, *Adamtsl2*-KD specific DEGs (n = 4,607 of 5,720) regulate bone and fat cell differentiation as well as growth factor signaling including WNT and BMP (Fig. 2K). Molecular Signature Database (MSigDB) analysis showed much higher enrichment of myogenesis-related genes in the shared and *Adamtsl2*-KD-specific DEGs compared to *Myod1*-KO-specific DEGs based on the combined score (Fig. 2L). This analysis raised the question of whether MYOD may execute of the myogenic differentiation program through regulating ADAMTSL2. We first analyzed the ADAMTSL2 promoter sequence upstream of the transcriptional start site and identified 13 and 10 putative MYOD binding sites in mouse and human, respectively, suggesting direct regulation of ADAMTSL2 by MYOD (Fig. 2M). Next, we asked if ADAMTSL2 could rescue lack of differentiation of MYOD-deficient C2C12 myoblasts. Knockdown of MYOD (*Myod1*-KD) using constitutive expression of *Myod1*- directed shRNA resulted in a ∼80% reduction of MYOD mRNA expression and a concomitant strong reduction of ADAMTSL2 mRNA supporting direct regulation of *Adamtsl2* by MYOD (Fig. 2N). Strikingly, overexpression of recombinant ADAMTSL2 or its C-terminal thrombospondin type 1 repeat (TSR) 2-4 domains (see below) not only fully rescued the differentiation defect of *Myod1*-KD myoblasts but even overcompensated resulting in a ∼2-fold increase in the fusion index compared to sh-Ctrl (Fig. 2O, P). Together, these data suggest that ADAMTSL2 as a critical effector of the MYOD differentiation program in myoblasts.

### ADAMTSL2 regulates canonical WNT signaling

To determine how ADAMTSL2 regulates myoblast differentiation downstream of MYOD, we first considered regulation of TGFβ signaling, since elevated TGFβ signaling due to ADAMTSL2 depletion would be consistent with reduced myoblast differentiation (Massague et al. 1986; Liu et al. 2001). We first verified the responsiveness of C2C12 myoblast to canonical TGFβ signaling, which induced SMAD2/3 phosphorylation (Fig. 3A). In contrast, SMAD2/3 phosphorylation was not changed in *Adamtsl2*-KD myoblasts, suggesting TGFβ-independent effects on myoblast differentiation. Next, we considered disruption of WNT signaling in *Adamtsl2*-KD myoblasts since (i) DEGs associated with WNT/β-catenin signaling were enriched in shared and *Adamtsl2-*KD-specific DEGs and (ii) WNT signaling is required for myoblast differentiation (Cossu and Borello 1999; Tanaka et al. 2011; Suzuki et al. 2015; Li et al. 2020). We used stabilization of β-catenin upon binding of canonical WNT ligands to their receptors and its translocation into the nucleus to measure canonical WNT/β-catenin signaling (Fig. 3B). Total β-catenin levels as determined by western blotting were reduced in differentiating *Adamtsl2*-KD myoblasts and increased when ADAMTSL2 was overexpressed or added as recombinant protein suggesting that ADAMTSL2 potentiates WNT/β-catenin signaling (Fig. 3C-E). Lithium chloride (LiCl), a known activator of canonical WNT signaling was used as positive control (Hedgepeth et al. 1997). Consistent with positive regulation of WNT signaling, β-catenin levels visualized by immunofluorescence microscopy were also reduced in *Adamtsl2*-KD myoblasts and no nuclear localization was observed (Fig. 3F). In contrast, ADAMTSL2 overexpression resulted in the almost exclusive localization of β-catenin in the nucleus. To correlate temporal regulation of ADAMTSL2, β-catenin and MyHC during myoblast differentiation, we analyzed cell lysates of differentiating myoblasts by western blotting. Consistent with ADAMTSL2 mRNA kinetics, ADAMTSL2 peaked at day 2 after differentiation initiation and correlated with a peak in β-catenin, followed by induction of MyHC (Fig. 3G). In lysates from *Adamtsl2*-KD myoblasts, ADAMTSL2 protein was significantly reduced and unchanged during myoblast differentiation (Fig. 3G). Concomitantly, cellular β- catenin levels not associated with WNT signaling remained constant at a low level and MyHC was not induced.

**Figure 3.**
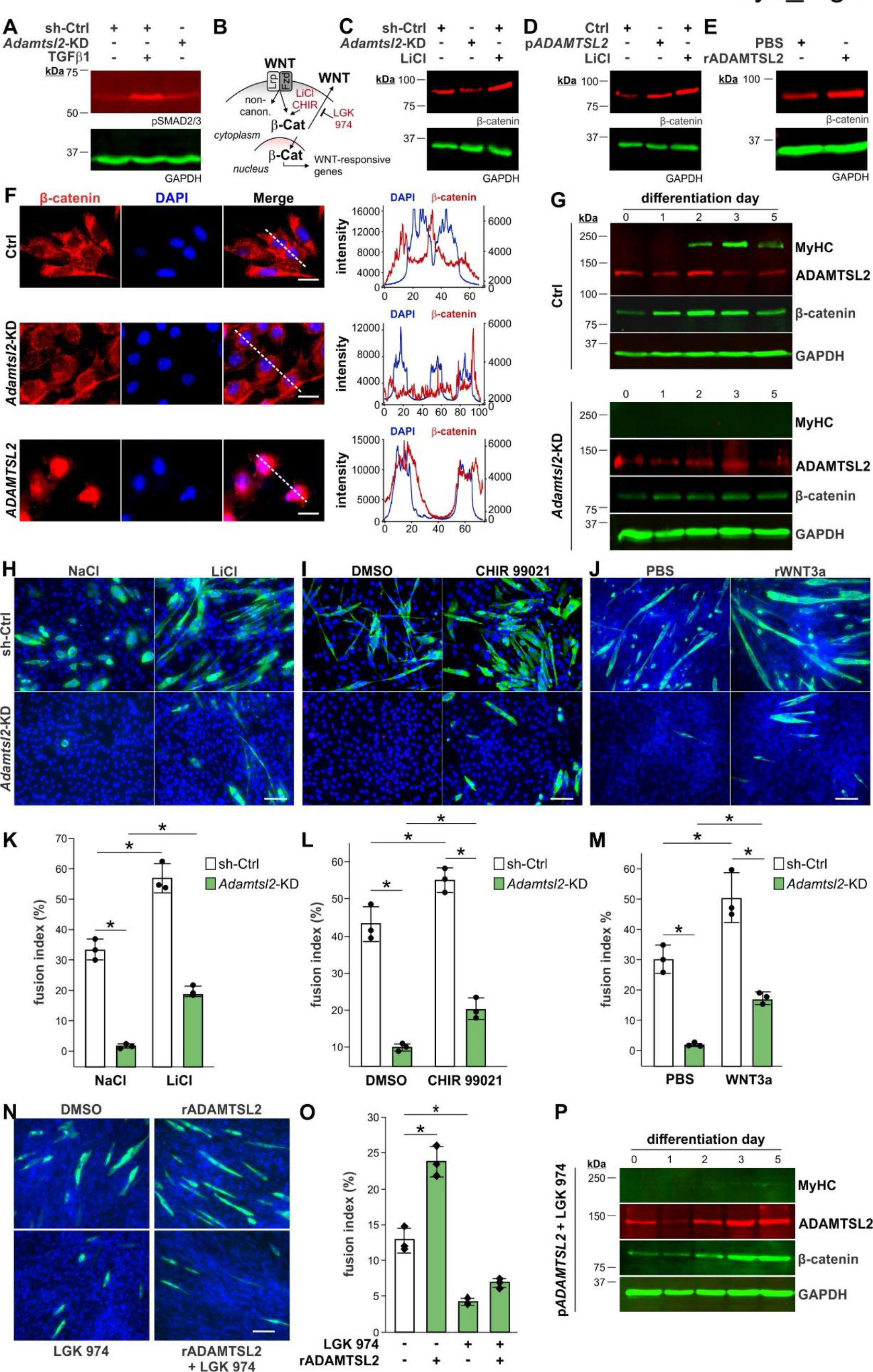
ADAMTSL2 promotes canonical WNT/β-catenin signaling. (*A*) Western blot analysis of SMAD2/3 phosphorylation in lysates from sh-Ctrl and *Adamtsl2-KD* C2C12 myoblasts under proliferating conditions. 10 ng/ml Recombinant TGFβ1 was used as positive control. (*B*) Schematic of canonical WNT signaling and targets of inhibitors. β-Cat, β-catenin. (*C*) Western blot analysis of β-catenin in sh-Ctrl and *Adamtsl2-KD* C2C12 myoblast lysates under proliferating conditions. (*D*) Western blot analysis of β-catenin in Ctrl and p*ADAMTSL2* overexpressing C2C12 myoblast lysates under proliferating conditions. LiCl was used as positive control in C and D. (*E*) Western blot analysis of β-catenin in C2C12 myoblast lysates treated with 100 µg/ml ADAMTSL2 protein (rADAMTSL2) under proliferating conditions. GAPDH was used as a loading control in A, C-E. (*F*) β-catenin immunostaining and intensity profiles along lines indicated in the merged image in proliferating Ctrl, *Adamtsl2-KD* and p*ADAMTSL2* overexpressing C2C12 myoblasts. (*G*) Western blot analysis of ADAMTSL2, MyHC, and β-catenin in sh-Ctrl (top) and *Adamtsl2-KD* (bottom) C2C12 myoblast lysates during differentiation. (*H-J*) MyHC immunostaining of sh-Ctrl and *Adamtsl2-KD* C2C12 myoblasts treated with 10 mM LiCl (H), 5 µM CHIR 99021 (I) or 100 ng/ml recombinant WNT3a (rWNT3a) (J). (*K-M*) Quantification of fusion indices from H-J (n = 3). (*N*) MyHC immunostaining of C2C12 myoblasts differentiated in the presence of 100 μg recombinant ADAMTSL2 and the porcupine inhibitor LGK 974 at 3 days after differentiation initiation. (*O*) Quantification of fusion index from N (n=3). (*P*) Western blot analysis of ADAMTSL2, MyHC, and β-catenin in differentiating p*ADAMTSL2* overexpressing C2C12 myoblast lysates treated with LGK 974. Scale bars: 20 μm (F), 100 μm in H-J. Bars represent mean ±SD. P-values were determined by one-way ANOVA and posthoc Tukey test. *p<0.05.

If ADAMTSL2 promotes myogenesis exclusively through regulation of canonical Wnt signaling, then pharmacological stabilization of β-catenin with LiCl or CHIR 99021 or addition of recombinant WNT3a ligand are predicted to rescue lack of differentiation of *Adamtsl2*-KD myoblasts (Fig. 3B) (Ring et al. 2003; Clement-Lacroix et al. 2005). When *Adamtsl2*-KD myoblasts were differentiated in the presence of LiCl, CHIR 99021 or Wnt3a, we observed significant rescue of myotube formation (Fig. 3H-J). However, the fusion index was not restored to control levels, suggesting the involvement of non-canonical WNT or other signaling pathways (Fig. 3K-M). To test if ADAMTSL2 required WNT ligands to regulate myoblast differentiation or if ADAMTSL2 itself could activate Wnt signaling, we differentiated C2C12 myoblasts in the presence of rADAMTSL2 and the porcupine inhibitor LGK 974, which blocks secretion of all WNT ligands (Barrott et al. 2011; Liu et al. 2013). As aforementioned, rADAMTSL2 promoted myoblast differentiation resulting in a ∼1.8-fold increase in the fusion index under control conditions (Fig. 3N, O). However, LGK 974 inhibited differentiation of C2C12 myoblasts in the presence of rADAMTSL2. In addition, myoblasts overexpressing ADAMTSL2 did not differentiate in the presence of LGK 974 and MyHC was not induced, despite robust ADAMTSL2 protein expression (Fig. 3P). Collectively, these data indicate that ADAMTSL2 promotes myogenesis through potentiation of canonical and possibly regulation of non-canonical WNT signaling.

### ADAMTSL2 binds to WNT ligands and WNT receptors

Since both ADAMTSL2 and WNT ligands are secreted by various muscle cell populations, we posited that ADAMTSL2 might potentiate WNT signaling by directly interacting with WNT ligands and/or WNT receptors. We investigated potential interactions using co-immunoprecipitation and molecular docking experiments. When Myc-tagged ADAMTSL2 was co-expressed with the extracellular domain of the canonical WNT co-receptor LRP6, ADAMTSL2 was co-immunoprecipitated by LRP6 in the presence or absence of WNT3a ligand (Fig. 4A). When Myc-tagged ADAMTSL2 was co-expressed with V5-tagged WNT3a, ADAMTSL2 was co-immunoprecipitated by WNT3a and vice versa (Fig. 4B, C). In addition to WNT3a, Myc-tagged ADAMTSL2 was also co-immunoprecipitated by V5-tagged WNT5a (non-canonical) and V5-tagged WNT7a (canonical/non-canonical), both WNT ligands are relevant for myogenesis during skeletal muscle formation or regeneration (Fig. 4D) (Le Grand et al. 2009; von Maltzahn et al. 2011; Reggio et al. 2020). To map the WNT-ligand binding sites in ADAMTSL2, we co-expressed the Myc-tagged N- and C-terminal ADAMTSL2 domains with V5-tagged WNT3a (Fig. 4E). By co-immunoprecipitation we localized one WNT3a binding site in the TRS1-Cys domains and one in the TSR2-7 domains (Fig. 4F). Based on equal input, WNT3a binding to TSR2-7 appeared to be much stronger.

**Figure 4.**
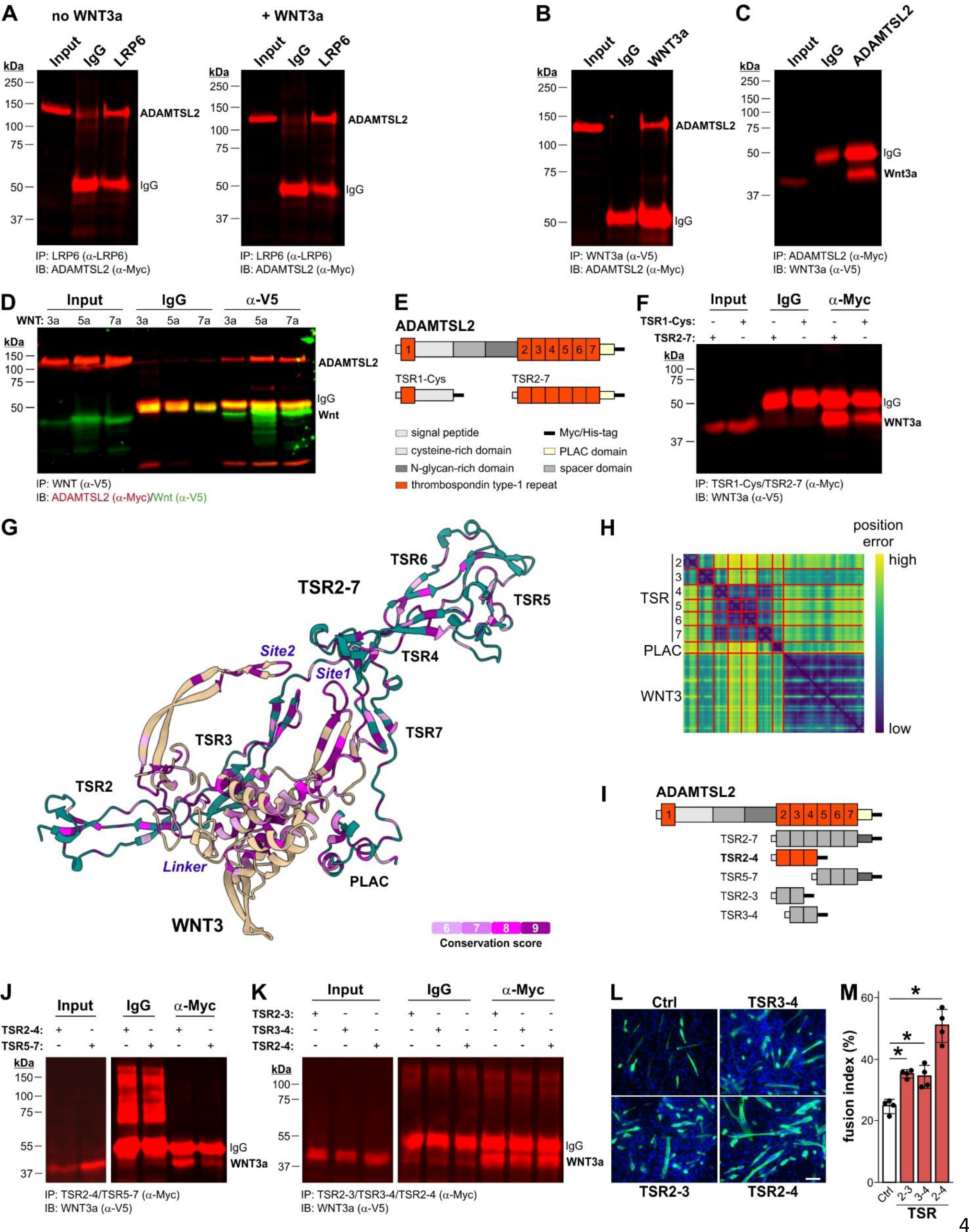
(*A*) Western blot analysis of ADAMTSL2 co-immunoprecipitated by LRP6 without (left) or with (right) WNT3a overexpression. (*B*) Western blot analysis of ADAMTSL2 co-immunoprecipitated by WNT3a. (*C*) Western blot analysis of WNT3a co-immunoprecipitated by ADAMTSL2. (*D*) Western blot analysis of ADAMTSL2 co-immunoprecipitated by WNT3a, WNT5a or WNT7a. (*E*) Domain organization of ADAMTSL2, TSR1-Cys and TSR2-7 constructs. (*F*) Western blot analysis of WNT3a co-immunoprecipitated by TSR1-Cys and TSR2-7. (G) AlphaFold modeling and prediction of WNT3 interactions with TSR2-7 domains of ADAMTSL2. Site 1 and 2 of WNT3 mediate the interaction with frizzled receptors and the WNT3 linker region mediates the interaction with LRP6. (*H*) Heat map of predicted local distance difference test (pLDDT) scores for G. (*I*) Domain organization of C-terminal ADAMTSL2 constructs. (*J*) Western blot of WNT3a co-immunoprecipitated by TSR2-4 or TSR5-7. (*K*) Western blot of WNT3a co-immunoprecipitated by TSR2-3, TSR3-4 or TSR2-4. (*L*) MyHC staining of C2C12 myoblasts transiently transfected with empty vector or plasmids encoding TSR2-3, TSR3-4 and TSR2-4 3 days after differentiation initiation. Scale bar: 100 µm. (*M*) Quantification of fusion index from (L) (n = 3). Bars represent mean ±SD. P-values were determined by one-way ANOVA and post-hoc Tukey test. *p<0.05. IP, immunoprecipitation; IB, immunoblot.

To gain further molecular insights, we used AlphaFold to model the interaction between the C-terminal TSR2-7 domains of human ADAMTSL2 and WNT3 based on their respective primary amino acid sequences. The AlphaFold model of the TSR2-7 – WNT3 complex placed the TSR2-4 domains of ADAMTSL2 adjacent to WNT3 with the greatest inter-domain interactions predicted to be with TSR3 and TSR4 (Fig. 4G). This is also reflected in the heat map of the predicted Local Distance Difference Test scores for the accuracy of the given alignment of individual TSR domains with WNT3 (Fig. 4H). To validate the AlphaFold model, we fine-mapped the WNT3a binding site in the C-terminal region of ADAMTSL2 by co-immunoprecipitation using constructs spanning the TSR2-7 domains (Fig. 4I). Consistent with the AlphaFold model, we localized the WNT3a binding site to TSR3 since Myc-tagged TSR2-3 and TSR3-4 co-immunoprecipitate V5-tagged WNT3a equally efficient (Fig. 4J, K). However, promotion of myoblast formation required the TSR2-4 domains, which caused a ∼2-fold increase in the fusion index after transient expression compared to a ∼1.4-fold increase for TSR2-3 or TSR 3-4 (Fig. 4L, M). Together, our data show that ADAMTSL2 may promote WNT signaling by directly binding to WNT ligands and WNT receptors.

### *Adamtsl2* deletion in myogenic precursor cells compromises skeletal muscle architecture

To determine the role of ADAMTSL2 in vivo, we first characterized the spatiotemporal expression pattern of *Adamtsl2* in skeletal muscle. *Adamtsl2* was expressed in tibialis anterior (TA), EDL, gastrocnemius (GM) and soleus (SOL) muscles with an up to 3-fold difference between individual muscles (Fig. 5A). *Adamtsl2* declined during postnatal growth in the TA muscle consistent with a largely developmental and early postnatal role for ADAMTSL2 in other tissues (Fig. 5B) (Hubmacher et al. 2015; Delhon et al. 2019; Hubmacher et al. 2019).

**Figure 5.**
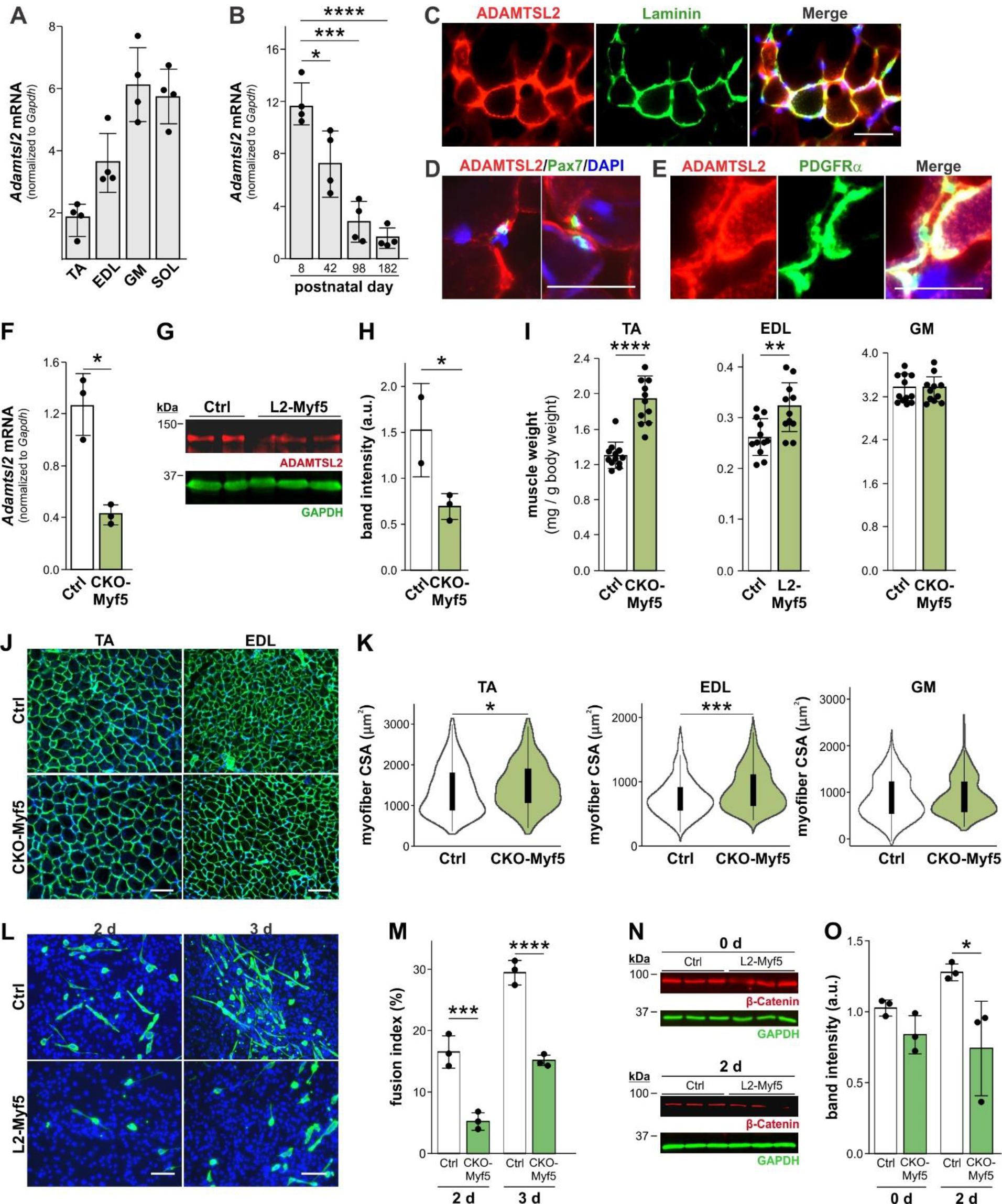
ADAMTSL2-depletion in *Myf5*-positive myogenic progenitor cells alters skeletal muscle architecture. (*A*) Normalized *Adamtsl2* mRNA levels in 4-week-old wild-type tibialis anterior (TA), extensor digitorum longus (EDL), gastrocnemius (GM) and soleus (SOL) muscle (n = 4). (*B*) Normalized *Adamtsl2* mRNA levels in wild-type TA muscle during postnatal growth (n = 4). (*C*) ADAMTSL2 and laminin immunostaining of wild-type TA cross-sections. (*D*) ADAMTSL2 and PAX7 immunostaining of wild-type TA cross-sections. (*E*) ADAMTSL2 and PDGFRα immunostaining of wild-type TA cross-sections. (*F*) *Adamtsl2* expression in TA muscle after *Myf5*-Cre-mediated *Adamtsl2* ablation (L2-Myf5) compared to Ctrl (n = 3). (*G*) Western blot analysis of ADAMTSL2 in extracts from Ctrl and L2-Myf5 EDL muscle. GAPDH was used as loading control. (*H*) Quantification of band intensity in B normalized to GAPDH. (*I*) TA, EDL and GM muscle weight from 8-week-old L2-Myf5 and Ctrl mice normalized to body weight (n = 11). (*J*) Laminin immunostaining of TA and EDL cross sections from 4-week-old L2-Myf5 and Ctrl mice. (*K*) Violin plots of myofiber cross-sectional area from J (n = 3). Boxes represent 25^th^-75^th^ percentile, whiskers ±S.D. (*L*) MyHC immunostaining of primary L2-Myf5 and Ctrl EDL-derived myoblasts at day 2 and 3 after initiation of differentiation. (*M*) Quantification of fusion index from (G) (n = 3). (*N)* Western blot analysis of β-catenin in protein extracts from Ctrl and L2-Myf5 primary myoblasts. GAPDH was used as loading control. (*O)* Quantification of band intensity from N normalized to GAPDH (n = 3). Scale bar in C, D, E: 50 µm; in J, L: 100 µm. Bars in A, B, F, H, I, M. O represent mean ±SD. P-values were determined with a Student’s t-test. *p<0.05, **p<0.01, ***p<0.001, ****p<0.0001.

Immunostaining of TA and EDL muscle cross-sections showed ADAMTSL2 localization in the endomysium surrounding myofibers (Fig. 5C, and Supplemental Figure S1*A, B*). ADAMTSL2 staining was strong near PAX7-positive satellite cells and PDGFRα-positive FAPs (Fig. 5D, E). To determine, if ADAMTSL2 is required for muscle development in vivo, we inactivated *Adamtsl2* in muscle precursor cells by combining a *Myf5*-Cre allele with a previously described conditional *Adamtsl2* allele (*Adamtsl2*^fl/fl^) resulting in CKO-Myf5 mice. *Adamtsl2* mRNA and protein were significantly reduced in 4-week old CKO-Myf5 TA muscle compared to *Adamtsl2*^fl/fl^ Ctrl littermates (Fig. 5F-H). We observed increased muscle weight and myofiber cross-sectional area in CKO-Myf5 TA and EDL, but not GM muscles (Fig. 5I-K). When *Adamtsl2* was inactivated in mature myofibers using muscle-specific creatinine kinase (*Ckmm*)-Cre, muscle weight and myofiber cross-sectional area did not change, indicating that ADAMTSL2 does not play a role in mature myofibers under homeostatic conditions (Supplemental Figure S1C-E). Consistent with our in vitro data from *Adamtsl2*-KD myoblasts, differentiation of primary myoblasts from Ctrl and CKO-Myf5 EDL muscles was delayed resulting in a ∼2-3-fold reduction in the fusion index at 2 and 3 days after initiation of differentiation (Fig. 5L, M). In addition, β-catenin levels were reduced in differentiating (day 2), but not proliferating (day 0) primary CKO-Myf5 myoblasts (Fig. 5N, O). These data suggest that ADAMTSL2 in myogenic-precursor cells is required for adequate muscle development and/or postnatal growth in a WNT-dependent manner.

## Discussion

Skeletal muscle formation is regulated through dynamic interactions between muscle-resident cells and their respective microenvironments as well as through secreted niche ligands. Key players in these regulatory networks are matricellular proteins, such as ADAMTSL2, which are ECM proteins that predominantly have regulatory rather than structural roles and are characterized by a highly dynamic and cell type-specific expression patterns (Murphy-Ullrich and Sage 2014). Here we identified an unexpected regulatory function for ADAMTSL2 in skeletal muscle as a regulator of myogenesis. Mechanistically, ADAMTSL2 promoted WNT signaling downstream of MYOD, a key myogenic transcription factor, by interacting with WNT ligands and the canonical WNT co-receptor LRP6. Therefore, ADAMTSL2 may facilitate the interaction between WNT ligands and their receptors or increase the effective concentration of WNT ligands at the cell surface or in the ECM.

ADAMTSL2 has previously been described as a negative regulator of TGFβ signaling in bronchial smooth muscle cells, skin and cardiac fibroblasts, and in the growth plate (Le Goff et al. 2008; Hubmacher et al. 2015; Delhon et al. 2019; Rypdal et al. 2021). Since lack of myoblast differentiation in the absence of ADAMTSL2 would have been consistent with elevated TGFβ signaling we first considered dysregulation of TGFβ-signaling in ADAMTSL2- deficient myoblasts (Massague et al. 1986; Liu et al. 2001). However, we did not detect alterations in canonical TGFβ signaling in ADAMTSL2-deficient myoblasts and it remains unclear if elevated TGFβ signaling is a driver of the phenotypes observed in skin fibroblasts, lung, heart, or cartilage or if TGFβ signaling increases as a consequence of an altered ECM structure . In addition, TGFβ signaling was not altered in mouse models for Weill-Marchesani syndrome, a connective tissue disorder that together with GD belongs to the acromelic dysplasia group and features the same musculoskeletal pathologies (Sengle et al. 2012; Mularczyk et al. 2018).

As an alternative to regulation of TGFβ signaling by ADAMTSL2, we considered regulation of canonical WNT signaling based on its prominent role in myoblast differentiation and skeletal muscle development and the enrichment of DEGs from *Adamtsl2*-KD myoblasts in WNT signaling-related molecular pathways (Girardi and Le Grand 2018). Canonical WNT signaling is required for myoblast differentiation at multiple steps on their trajectory towards myotubes (Cossu and Borello 1999; Brack et al. 2008; Tanaka et al. 2011; Suzuki et al. 2015). Using a porcupine inhibitor, which blocks secretion of all WNT ligands, we abolished the capacity of ADAMTSL2 to promote myoblast differentiation. This dependency of the pro-myogenic function of ADAMTSL2 on secreted WNT ligands strongly suggested that it is mediated through canonical and/or non-canonical WNT signaling. It also showed that ADAMTSL2 did not activate Wnt signaling independent of WNT ligands. Regulation of canonical WNT signaling was further supported by the significant rescue of differentiation of ADAMTSL2-deficient myoblasts. However, the fact that we only partially rescued ADAMTSL2-deficiency suggested that ADAMTSL2 could potentially regulate both, canonical and non-canonical WNT signaling.

This possibility was supported by our findings that ADAMTSL2 could bind to several, possibly all canonical and non-canonical all WNT ligands. In vivo, ADAMTSL2 may be agnostic to individual WNT ligands and consequently may modulate WNT signaling solely depending on the WNT ligands that are present in a given microenvironment. Alternatively, ADAMTSL2 could shift the balance between canonical and non-canonical WNT signaling or integrate WNT signaling with other signaling pathways, such as TGFβ signaling (Carthy et al. 2011; Biressi et al. 2014; Dzialo et al. 2018; Contreras et al. 2020). Detailed competition experiments with opposing WNT ligands in systems that are responsive to both, canonical and non-canonical WNT signaling, could address this question. Our biochemical and structural data further suggest that ADAMTSL2 could promote the WNT ligand/WNT receptor interaction and by doing so potentiate WNT signaling. It is possible that ADAMTSL2 forms a complex with the WNT ligand, the LRP6, and even the Frizzled receptor and thus sequesters all three WNT signaling components in close proximity to facilitate their interaction or to increases the affinity for WNT ligands to their receptors (Hirai et al. 2019). In the predicted AlphaFold interaction model of the C-terminus of ADAMTSL2 and WNT3, the sites that bind to frizzled- and LRP6- co-receptors are predicted to be accessible (Hirai et al. 2019). As a precedent, it was shown that the ECM protein biglycan bound to WNT3a and promoted its activity by simultaneously binding to LRP6 (Berendsen et al. 2011). Several secreted molecules, such as secreted Frizzled-related proteins (sFRPs), dickkopf (Dkk), or R-spondins can modulate WNT signaling (Cruciat and Niehrs 2013). However, most of these molecules repress WNT signaling, with the exception of R-spondins and Norrin (Xu et al. 2004; Kim et al. 2008). In this regard, ADAMTSL2 behaves more like an R-spondin, since its WNT-modulatory activity was dependent on the presence of WNT ligand(s). In contrast to R-spondins however, ADAMTSL2 directly bound WNT ligands, while R-spondins stabilized the LRP6 receptor at the cell surface (Zebisch and Jones 2015; Dubey et al. 2020). ADAMTSL2 likely functions as a rheostat rather than an on-off switch for WNT signaling and myogenesis. Thus, myoblasts can fine-tune WNT signaling through transcriptional regulation of ADAMTSL2 without the need to regulate individual components of the WNT signaling pathway.

One surprising finding was that ADAMTSL2 overexpression fully rescued lack of differentiation of MYOD-deficient myoblasts. This suggested that ADAMTSL2 is a key effector molecule downstream of MYOD, which was corroborated by the identification of multiple putative MYOD binding sites in the promoter region of the human and mouse ADAMTSL2 gene. In addition, DEGs of differentiating *Adamtsl2*-KD and *Myod1*-KO myoblasts substantially overlapped and the overlapping DEGs were enriched in pathways related to myogenesis and skeletal muscle biology. This indicates that ADAMTSL2 may regulate the same myogenic pathways as MYOD, which as a transcription factor regulates transcription of target genes. Cellular changes however, are driven by effector molecules downstream of MYOD. As such, ADAMTSL2 could be key mediator a MYOD-ADAMTSL2-WNT axis that promotes myoblast differentiation during muscle formation. One limitation is that the consequences of MYOD ablation in vivo are not fully recapitulated by MYOD ablation in C2C12 myoblasts. *Myod1* KO mice have no developmental muscle phenotype due to compensation by the myogenic regulatory factor MYF5 (Rudnicki et al. 1992). Only when both, *Myod1* and *Myf5* were knocked out myofibers were not formed (Rudnicki et al. 1993). We observed a similar behavior, where knockdown of ADAMTSL2 in C2C12 myoblasts resulted in complete lack of differentiation, while differentiation of primary ADAMTSL2-deficient myoblasts was delayed, but not abolished.

Elucidating the reasons for this discrepancy and the question what can potentially compensate for lack of ADAMTSL2 in myogenic precursor cells in vivo needs further investigation.

Pathogenic variants in *ADAMTSL2* cause GD in humans resulting in short stature and a pseudomuscular build characterized by apparent muscle hypertrophy (Le Goff et al. 2008). In congruence, inactivation of *Adamtsl2* in myogenic precursor cells in vivo resulted in increased muscle mass and myofiber cross-sectional area. This would suggest that dysfunctional or absent ADAMTSL2 enhances myogenesis or promotes muscle fiber hypertrophy. However, our gain- and loss-of-ADAMTSL2 function studies in myoblasts in vitro suggested that ADAMTSL2 promotes myogenesis and the prediction for a loss-of-function phenotype in vivo would be reduced muscle size or impaired myogenesis. How can these contrasting findings be reconciled? First, C2C12 myoblasts were derived from adult myoblasts after injury (Yaffe and Saxel 1977). Therefore, C2C12 cells are more representative of adult skeletal muscle regeneration after injury rather than embryonic or postnatal skeletal muscle development, which differs with respect to myoblast origin and growth factor sensitivities (Hutcheson et al. 2009). Second, most *ADAMTSL2* variants causing GD are single nucleotide variants and so far no individual with GD with a complete lack of *ADAMTSL2* has been reported. The pathogenic *ADAMTSL2* variants that were analyzed biochemically resulted in reduced, but not abolished secretion (Bader et al. 2010; Piccolo et al. 2019). This would suggest that residual amounts of ADAMTSL2 are secreted in GD and a complete knockout of *Adamtsl2* may not recapitulate the consequence of GD-causing pathogenic variants in ADAMTSL2, including potential negative gain of functions. This is supported by the reported almost 100% perinatal lethality of *Adamtsl2* KO mice, while close to 70% of patients with GD survive childhood (Hubmacher et al. 2015; Marzin et al. 2020). In addition, GD-causing pathomechanisms could be related to the accumulation of mutant ADAMTSL2 protein within the secretory pathway, which may compromise autophagy, induce the unfolded protein response, or result in aberrant activation of TGFβ signaling (Piccolo et al. 2019). To distinguish between consequences of lack of ADAMTSL2 in *Adamtsl2* KO mice and that of GD-causing *ADAMTSL2* variants, knock-in mice harboring individual GD-causing pathogenic *Adamtsl2* variants will be required.

In summary, we identified a novel function for ADAMTSL2 in regulating WNT signaling during myogenesis. It will be exciting to elucidate further the role of ADAMTSL2 in potentially mediating tissue-specific WNT-TGFβ signaling crosstalk and to determine the role of different pools of ADAMTSL2 in skeletal muscle contributed by differentiating myogenic precursor cells and muscle connective tissue cells during skeletal muscle development and regeneration.

Finally, it will be important to define the pathogenic mechanisms of GD and to examine the consequences of disease-causing variants in *ADAMTSL2*, not only in skeletal muscle, but also during development and homeostasis of other tissues affected in GD patients, such as the trachea, the lungs, the skin, or the heart valves.

## Materials and Methods

### Mice

B6.129S4-*Myf5^tm3(cre)Sor^*/J (JAX 007893) and B6.FVB(129S4)-Tg(*Ckmm*-cre)^5Khn^/J (JAX 006475) strains were purchased from the Jackson Laboratory (Bar Harbor, ME) (Bruning et al. 1998; Tallquist et al. 2000). Mice harboring the conditional *Adamtsl2* allele were described previously (Hubmacher et al. 2019). Mice were used with approval from the Institutional Animal Care and Use Committee at the Icahn School of Medicine at Mount Sinai (IACUC-2018-0009, PROTO202000259). Mice were housed in a temperature-controlled environment with free access to food and water under a 12 h light/dark cycle. Mice were euthanized by CO2 asphyxiation followed by cervical dislocation prior to tissue or cell extraction

### Cell culture

Human embryonic kidney (HEK) 293 cells (CRL-1573, ATCC) and C2C12 myoblasts (CRL- 1772, ATCC) were cultured in Dulbecco’s modified Eagle’s medium (DMEM, GIBCO) supplemented with 10% fetal bovine serum, 1 mM sodium pyruvate (GIBCO), 100 units penicillin/100 μg/ml streptomycin (GIBCO) (complete DMEM) (Yaffe and Saxel 1977). C2C12 myoblasts were maintained by passaging at low cell densities for up to 25 passages. Differentiation was induced by switching confluent C2C12 myoblasts to DMEM supplemented with 2% horse serum (GIBCO), 1 mM sodium pyruvate and 100 units penicillin/100 μg/ml streptomycin (differentiation medium). Cultured cells were incubated in a humidified incubator in a 5% CO2 atmosphere at 37 °C.

### Isolation of primary myogenic progenitor cells (myoblasts)

EDL muscles from wild type mice were minced and digested with 1 ml 0.2% Collagenase IV (Worthington) in DMEM with 1 mM sodium pyruvate at 37 °C for 45 min to release individual myofibers (Pasut et al. 2013). Myofibers were transferred into one well of a 6-well plate pre-coated with horse serum and incubated in DMEM with sodium pyruvate. This step was repeated twice resulting in the release of muscle progenitor cells from myofibers. Myoblasts were cultured in Ham’s F12 medium (GIBCO) supplemented with 1 mM sodium pyruvate, 20% fetal bovine serum, 0.5 nM FGF-2 (Sigma Aldrich), 100 units penicillin/100 μg/ml streptomycin. Differentiation of primary myoblasts at confluency was initiated by switching to differentiation medium.

### Transfection and generation of stable cell lines

Transfection of HEK 293 cells with plasmid DNA was performed with PEI reagent (Polysciences) in a 1:1 (w/w) ratio to plasmid DNA in serum-free Opti-minimal essential medium (Opti-MEM) (GIBCO). C2C12 myoblasts were transfected using TransfeX reagent (ATCC) at a ratio of 1 µg plasmid DNA to 2 µl transfection reagent according to the manufacturer’s instructions. For the generation of C2C12 myoblasts stably expressing human ADAMTSL2, we used lentivirus-mediated transduction of an expression plasmid (synthesized by Vector Builder), which contained a separate green-fluorescent protein (GFP) expression cassette for fluorescence-activated cell sorting (FACS). To produce the lentivirus, HEK 293T packaging cells (3 x 10^6^) were seeded in complete DMEM. After 24 h cells were co-transfected with 1.3 pmol of psPAX2 (Addgene, 12260), 0.72 pmol pCMV-VSV-G (Addgene, 8454) and 1.64 pmol ADAMTSL2 plasmids using PEI reagent. Supernatant containing the virus was harvested at 48, 72 and 96 h post transfection, centrifuged at 500 g for 5 min, passed through a 0.45 µm syringe filter, and stored in batches at -80 °C. To stably deplete ADAMTSL2 mRNA, C2C12 myoblasts were transfected with plasmids encoding small hairpin (sh)-RNA targeting *Adamtsl2* (Mission shRNA, Sigma Aldrich) using PEI. Stable cells were selected with 5 µg/ml puromycin (VWR) in complete DMEM for 3 days and subsequently maintained in complete DMEM plus 3 µg/ml puromycin. Knockdown efficiency of three individual shRNAs targeting different regions of the *Adamtsl2* mRNA (XM_130065.5-1977s1c1, XM_130065.5-3086s1c1, and XM_130065.5-972s1c1) was similar and we used XM_130065.5-3086s1c1 targeting the 3’-untranslated region (UTR) of *Adamtsl2*, because it would not interfere with rescue experiments using plasmids encoding recombinant ADAMTSL2.

### Cloning of plasmids encoding recombinant ADAMTSL2 domains

Recombinant human ADAMTSL2 constructs where generated by PCR-amplification with the Q5 Hot start high fidelity 2x master mix (NEB) and specific primer pairs to allow for restriction cloning. Cloning primers are listed in Supplemental Table S1. PCR products were amplified with the following program: 98 °C for 30 sec, 34 cycles of 98 °C for 10 sec, 55 °C for 30 sec and 72 °C for 30 sec/kb followed by a final extension for 2 min at 72 °C. The PCR products and the pSecTag 2B vector (ThermoFisher Scientific), which contains a signal peptide for secretion, were restriction-digested with BamHI × XhoI (NEB) and the agarose gel-purified DNA fragments were ligated with T4 DNA ligase (NEB) at 16 °C overnight. The N-terminal domains of ADAMTSL2, which include the endogenous signal peptide, were cloned in pcDNA3.1(-) Myc/His vector (ThermoFisher Scientific). Ligated plasmids were transformed in chemically competent *E. coli* DH5α cells (NEB, C2988J) and positive clones were identified by ampicillin resistance and verified by DNA sequencing.

### Immunoprecipitation

ADAMTSL2 and WNT3a, WNT5a, and WNT7a protein interaction studies were performed by co-transfecting HEK 293 cells in a 10 cm cell culture dish with 5 µg of pcDNA-WNT3a-V5, pcDNA-WNT5a-V5 or pcDNA-WNT7a-V5 (Addgene, 35927, 35930, 35933) and 5 µg of plasmids encoding full length ADAMTSL2 or Myc/His-tagged ADAMTSL2 peptides. For LRP6 and ADAMTSL2 protein interaction studies the LRP6-pCS2 (Addgene, 27242) was used for co-transfection. The WNT-V5-tagged plasmids were a gift from Marian Waterman and the LRP6-pCS2 was a gift from Xi He and described previously (Tamai et al. 2000; Najdi et al. 2012). Transfection was performed with PEI as described above. Transfected HEK 293 cells were switched to serum-free DMEM 24 h after transfection and cultured for an additional 2 days. Cell lysates were prepared using 1 ml RIPA lysis and extraction buffer (ThermoFisher Scientific), supplemented with cOmplete EDTA-free protease inhibitor cocktail (Roche) on ice for 30 min. Protein lysates were cleared by centrifugation at 12,000 g for 20 min at 4 °C. The protein concentration in the lysate was determined using a detergent-compatible Bradford assay (ThermoFisher Scientific). Co-immunoprecipitation was performed with 1 mg of protein from whole cell lysate or 4 ml of conditioned medium. Proteins binding non-specifically to protein A/G were removed by pre-adsorption of cell lysate or conditioned medium to 20 µl protein A/G magnetic beads (Pierce) for 1 h rotating at 10 rpm at 4 °C. Beads were pelleted with a magnet and protein lysates or conditioned medium were transferred into new Eppendorf tubes. 2 µg of antibodies against the V5-tag (Invitrogen, R960-25), the c-Myc tag (Invitrogen, clone 9E10, MA1-980), or LRP6 (Cell Signaling, clone C5C7, 2560) were added and incubated overnight at 4 °C under rotation. Mouse IgG (Genetex, #GTX35009) or rabbit IgG (Genetex, #GTX35035) were used as negative controls, Immunoprecipitated proteins were captured by adding 25 µl of magnetic A/G beads for 4 h at 4 °C under rotation. To remove unbound proteins, beads were pelleted and washed 3 × 5 min with 500 µl of RIPA lysis buffer under rotation at room temperature (RT). Proteins were eluted by adding 50 µl of 2x reducing SDS loading buffer, vortexing for 5 min, and incubation at 95 °C for 5 min. The eluted proteins were separated from the magnetic beads and subjected to SDS polyacrylamide gel electrophoresis (SDS-PAGE) followed by western blotting.

### SDS-PAGE and western blotting

SDS-PAGE was performed with 50 µg of proteins from cell lysates or from trichloroacetic acid (TCA)-precipitated conditioned medium. For TCA precipitation, 1 ml of conditioned medium was combined with 395 μl of premixed TCA (256 µl VWR Life Science) and 1% Triton-X 100 (139 µl, Acros Organic), vortexed for 30 sec and incubated on ice for 10 min. Precipitated proteins were pelleted by centrifugation at 10,000 rpm for 10 min at 4 °C. The protein pellet was rinsed twice with chilled acetone followed by centrifugation for 10 min at 4 °C and air dried for 5 min. The protein pellet was dissolved in 1x reducing SDS-loading buffer and incubated at 95 °C for 5 min. Proteins were resolved using SDS-PAGE and transferred onto a activated polyvinylidene difluoride (PVDF) membrane (EMD Millipore) at 400 mA for 3 hours using a MiniTransblot Module (BioRad). After protein transfer, membranes were blocked for 2 h with 5% milk in PBS including 0.1% Tween-20 (PBST) at RT or overnight at 4 °C. Membranes were then washed with PBST for 5 min and incubated with the following primary antibodies: GAPDH (EMD Millipore, #MAB 374, 1:2000), β-catenin (EMD Millipore, #05-613, 1:2000), LRP6 (Cell Signaling, clone C5C7, 2560), c-Myc (Invitrogen, clone 9E10, MA1-980, 1:1000), V5 (Invitrogen, #46-0705, 1:2000), ADAMTSL2 or myosin heavy chain (MyHC) (DSHB, #MF-20c, 1:1000) diluted in PBST for 2 h at RT. After washing the membranes 3 × 5 min with PBST, secondary antibodies goat anti-mouse (LI-COR, #IRDye 800CW/680RD) and goat anti-rabbit (LI-COR, #IRDye 680RD/800CW) at 1:10,000 dilution were added and incubated for 1 h at RT. Membranes were washed 3 × 5 min with PBST, 1 × 5 min with PBS and imaged using the Azure Biosystem c600 (Azure Biosystems). Band intensities of western blots were quantified with ImageJ. The ADAMTSL2 antibody was raised in rabbits against a peptide spanning the amino acids 552-567 of mouse ADAMTSL2 (CTHKARTRPKARKQGVS (YenZym Antibodies, LLC). The antibody was affinity purified using the immobilized immunogenic peptide.

### mRNA isolation, cDNA preparation, and quantitative real-time PCR

mRNA was extracted from cell layers and skeletal muscle tissues using TRIzol (ThermoFisher Scientific). Briefly, cells were lysed in 1 ml Trizol after removing the cell culture medium and mRNA was isolated according to the manufacturer’s protocol. To isolate mRNA from mouse skeletal muscle, individual muscles were dissected, snap-frozen in liquid nitrogen, and pulverized using a Geno/Grinder (Spex SamplePrep). Pulverized tissues was re-suspended in TRIzol and mRNA was isolated according to the manufacturer’s protocol. After air-drying, the pellet was dissolved in nuclease-free water and mRNA concentration and purity was determined with a Nanodrop spectrophotometer (Thermo Scientific). 1 µg of mRNA was digested with DNaseI (Thermo Scientific) prior to cDNA synthesis to remove traces of genomic DNA and reverse transcribed into cDNA using the High Capacity cDNA Reverse Transcription kit (Applied Biosystem) according to the manufacturer’s instructions. For quantitative real-time PCR analysis, cDNA was diluted (1:5) in nuclease-free water and a SYBR green PCR master mix (Applied Biosystem) was used to amplify gene specific PCR products with the primers listed in Supplemental Table S2. The PCR products were amplified using the following steps: 48 °C for 30 min, 95 °C for 10 min and 40 cycles of 95 °C for 15 sec and 60 °C for 1 min. The cycle threshold (ct) value was used to calculate the fold change using the 2^−(ΔΔCT)^ method and graphs were plotted using Origin 2019 (OriginLab).

### RNA sequencing

sh-ctrl and *Adamtsl2*-KD C2C12 myoblasts (3 × 10^5^/well) were seeded in triplicates in 6-well plates and cultured in complete DMEM until confluency. To induce differentiation, C2C12 myoblasts were cultured in differentiation medium for an additional 3 days with daily medium change. After 3 days, cells were lysed with 1 ml Trizol and mRNA was extracted according to the manufacturer’s protocol. RNA quality control, library preparation, sequencing and bioinformatics determination of DEGs was performed by Azenta Life sciences (Genewiz). mRNA quality was controlled using NanoDrop, RNA Qubit and Tape Station instruments.

Sequencing libraries were prepared by polyA selection for mRNA and sequenced to a depth of 20-30 Mio reads (HiSeq 4000 2 × 150 paired-end configuration, Illumina). Sequence reads were trimmed to remove possible adapter sequences and nucleotides with poor quality using Trimmomatic v.0.36. The trimmed reads were mapped to the *Mus musculus* GRCm38 reference genome (ENSEMBL) using STAR aligner v.2.5.2b. Unique gene hit counts were calculated by using featureCounts from the Subread package v.1.5.2. Only unique reads that fell within exon regions were counted. Gene hit counts were used for downstream differential expression analysis. Using DESeq2, a comparison of gene expression between the two groups of samples was performed. The Wald test was used to generate p-values and log2-fold changes. Genes with an adjusted p-value <0.05 and absolute log2-fold change >1 were called as differentially expressed genes for each comparison. Data are available at the GEO database (GSE185894)

### Immunofluorescence staining of cells

For immunofluorescence staining of proliferating and differentiating C2C12 myoblasts, 50,000 cells per chamber were seeded in 8-well glass chamber slides and cultured in complete DMEM until confluency. Differentiation was initiated by switching to differentiation medium. At the end of the culture period, cells were rinsed three times with PBS and fixed with ice-cold methanol (Fisher Chemicals) for 15 min at -20 °C. After three washes with PBS, cells were blocked with 5 % BSA in PBST for 2 h at RT. Cells were incubated with antibodies against β-catenin (EMD Millipore, # 05-613), Ki67 (Abcam, #ab16667), MyHC (DSHB, #MF 20), laminin (Sigma, #L9393 and Novus Bio, #NB600-883) or ADAMTSL2 diluted at 1:200 in PBS overnight at 4 °C. Cells were then washed 3 × 5 min with PBS and incubated with fluorophore-tagged goat-anti mouse or goat-anti rabbit secondary antibodies (Jackson Immuno Research, #Rhodamine Red 111-295-144 or 115-295-146 and Alexa Fluor 488 115-545-146 or 111-545-144) diluted at 1:500 in PBST for 1 h at RT. After washing 3 × 5 min with PBS, cells were mounted with DAPI containing mounting medium (Invitrogen) and observed using an Axio Imager Z1 fluorescence microscope (Zeiss).

### Immunofluorescence staining of muscle tissue

EDL, TA and GM muscles were dissected and partially embedded in a plastic cassette holder using 1 g/ml of tragacanth powder gum (Alfa Aesar) in water. Muscles where then immersed in liquid nitrogen-chilled 2-methylbutane (Fisher Chemicals) for 2 min. Subsequently, the fixed tissues were kept in liquid nitrogen for 1 min and wrapped in aluminum foil and stored at -80 °C. To prepare frozen sections, muscle tissues were embedded with optimal cutting temperature (OCT) medium (Tissue-Tek) and kept at -20 °C. 20 µm sections were obtained using a cryostat (Avantik) and mounted onto frosted glass slide. Sections were incubated with 4% PFA (MP) for 20 min at RT and washed 2 × 10 min with PBS. The tissue was permeabilized with chilled methanol for 6 min at -20 °C and washed for 10 min with PBS. 0.01

M citric acid (Fisher BioReagents) solution in water was heated by microwaving until the temperature reached 90 °C. Sections were immersed in the hot citric acid buffer and steamed for 5 min in the microwave. Sections were cooled down in citric acid for 30 min at RT and washed for 15 min with PBS. Sections were incubated with 5 % BSA in PBS for 2 h. Sections were then incubated in primary antibodies against laminin (LAM-89, NB600-883, Novus Biologicals), ADAMTSL2, MyHC, PAX7 (DSHB), PDGFRα (Invitrogen, #14-1401-82) were diluted 1:200 in PBS for overnight at 4 °C. Sections were washed 3 × 10 min with PBS and incubated with corresponding fluorophore-tagged secondary antibodies at 1:500 dilution for 1 h at RT. Slides were washed 3 × 10 min with PBS and mounted with DAPI-containing mounting medium.

### Prediction of the ADAMTSL2-WNT complex using AlphaFold

For the generation of ADAMTSL2 TSR2-7 and WNT3 models, AlphaFold multimer models were generated using a locally installed version of AlphaFold v2.1.1 on the Computationally Shared Facility at University of Manchester (Jumper et al. 2021). The installation included the full genetic databases for multiple sequence alignments. Sequences of ADAMTSL2 (Q86TH1: 564-951) and WNT3 (P56703-1:42-355) were used for modelling. FASTA files containing complexes of the TSR domains of ADAMTSL2 and WNT3 were prepared and submitted to the AlphaFold pipeline and ranked by the software. Models were visualized in UCSF ChimeraX (Pettersen et al. 2021).

### Statistical Analysis

Two sample comparisons were done with a two-sample Student’s t-test and multi-sample comparisons with a one-way ANOVA followed by a posthoc Tukey analysis to determine which samples were different. A p-value cutoff of <0.05 was considered significant. Sample sizes are indicated in the figure legends. OriginPro software was used for statistical calculations and plotting.

### Data Availability

RNA sequencing data were deposited in the GEO database (GSE185894). All other data are included in the article or in supplemental materials.

## Acknowledgments

Funding was provided by the National Institutes of Health grant R01AR070748 (D.H.), Biotechnology and Biological Sciences Research Council, UK, studentship (C.B.), Biotechnology and Biological Sciences Research Council, UK, grant BB/R008221/1 (C.B.), Wellcome Trust, UK, grant 203128/Z/16/Z (C.B., through Wellcome Centre for Cell-Matrix Research).

## Author Contributions

N.T. and D.H. designed research; N.T., M.S., and C.B. performed research; N.T., M.S., C.B., and D.H. analyzed data; N.T. and D.H. wrote manuscript and prepared the figures. All authors edited and approved final manuscript.

---

**Supplementary Figure S1:**
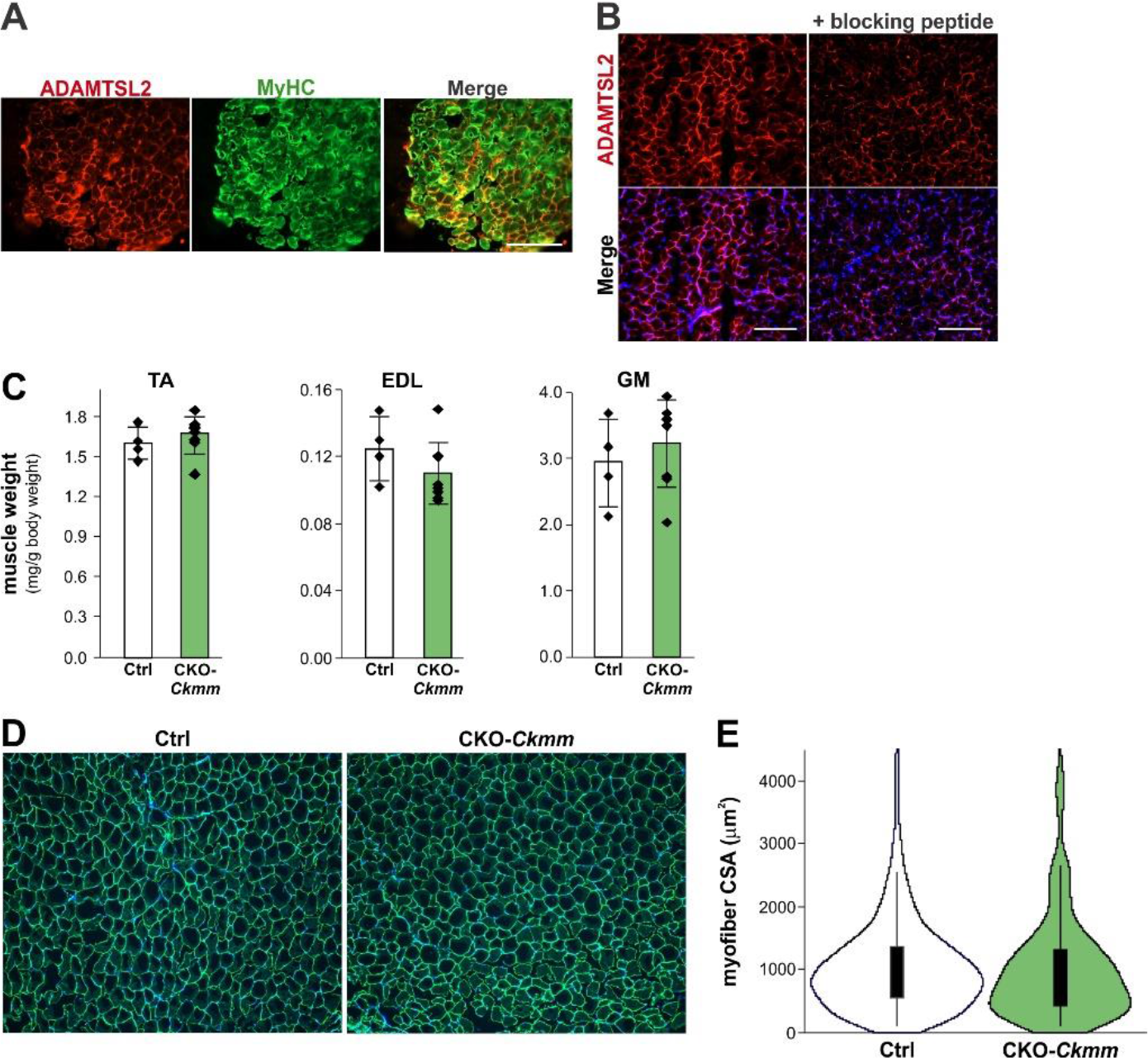
(*A*) ADAMTSL2 and MyHC co-immunostaining in cryosections of wild-type TA muscle at 4 weeks of age. (*B*) ADAMTSL2 immunostaining in cryosections of wild-type TA muscle at 4 weeks of age with or without blocking peptide. Representative images from n = 3 cryosections are shown. Scale bar: 100 µm. (*C*) Wet weight of TA, EDL and GM muscle dissected from L2-Ckmm and *Adamtsl2*^fl/fl^ controls normalized to the body weight at 8 weeks of age (n = 4-7). Bars represent mean ±SD. P-values were determined using a two-sided Student’s t-test and a p-value <0.05 was considered significant. (*D*) Laminin immunostaining of cryosections of TA and EDL muscle from L2-Ckmm and *Adamtsl2*^fl/fl^ controls at 8 weeks of age. (*E*) Violin blots of myofiber cross-sectional area from (*D*) (n = 3).

**Supplementary Table S1.**
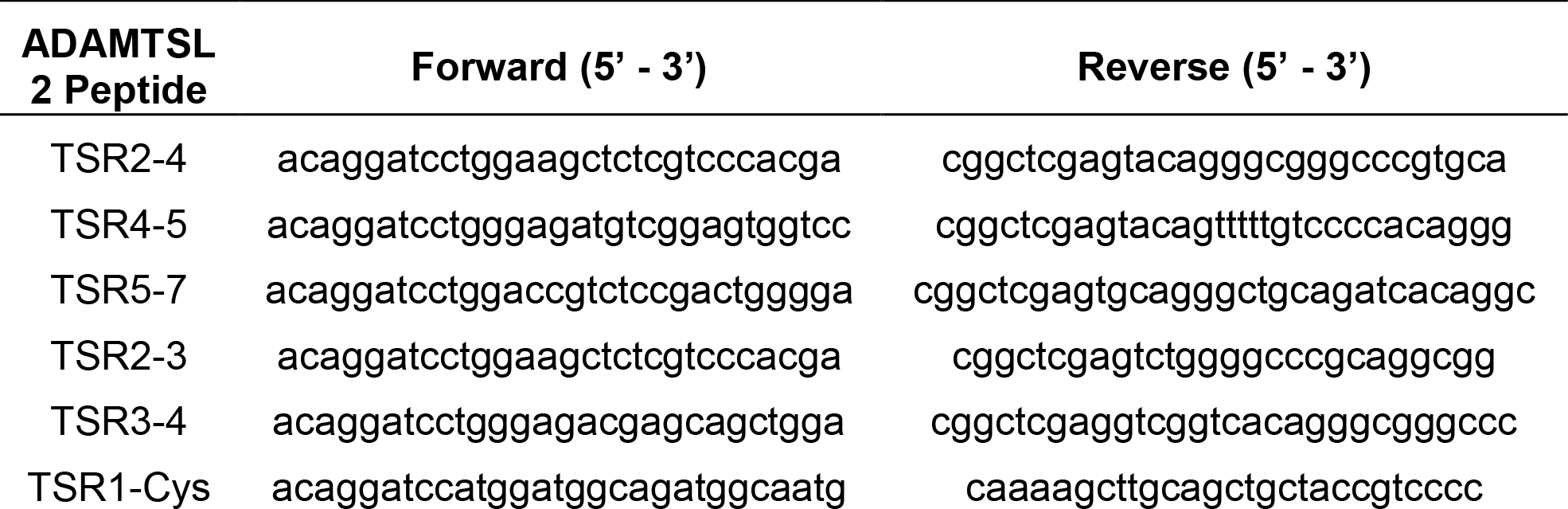
ADAMTSL2 cloning primer sequences

**Supplementary Table S2.**
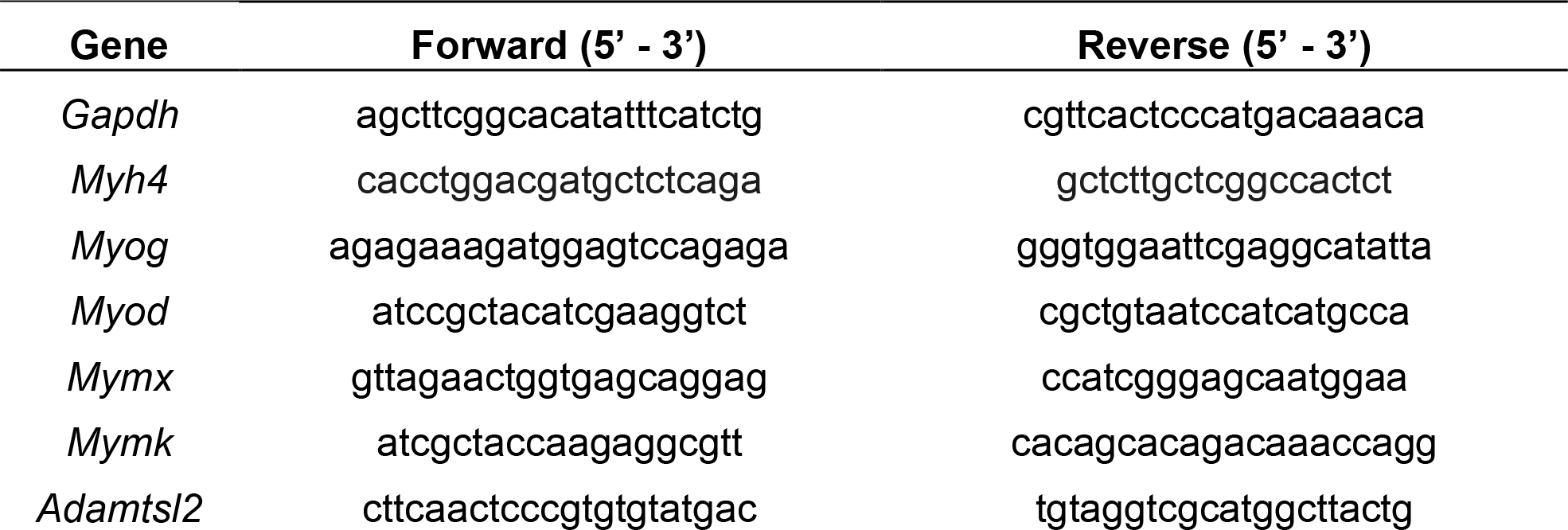
ADAMTSL2 qPCR primer sequences

